# Rational Molecular Design of Two-Photon Activated Temoporfin: A Computational Study for Advanced Photodynamic Therapy

**DOI:** 10.1101/2025.05.02.651861

**Authors:** Basak Koca Fındık, Ege Su Uyar, Antonio Monari, Saron Catak

## Abstract

Photodynamic therapy (PDT) is a promising, non-invasive cancer treatment that relies on the activation of photosensitizers (PS) by suitable light to produce cytotoxic reactive oxygen species. However, the efficiency of PDT is often hindered by the limited penetration of visible light into tissues, requiring the use of infrared activable PS. Furthermore, PS are usually prone to aggregation and present solubility issues limiting their bioavailability. In this study, we explore the functionalization of temoporfin (mTHPC), a clinically approved second-generation PS, with two-photon absorption (TPA) chromophores to enhance its efficiency in deep tissues. Three TPA-temoporfin conjugates (DTP1-mTHPC, DTP2-mTHPC, and DPP-mTHPC) have been designed and their properties have been investigated using a combination of quantum mechanics (QM), molecular dynamics (MD), and hybrid QM/MM simulations. Computational analysis revealed that the TPA cross-section (σ) of the parent DTP moieties significantly increase when anchored to mTHPC, thus allowing efficient absorption in the near-infrared (NIR) region. Additionally, we have shown that their encapsulation with β-cyclodextrins (β-CDs) improved solubility and prevented aggregation without altering the optical properties of the PS. Simulations in a biological membrane model confirmed favorable interactions and localization of the candidate PDT agents within lipid bilayers, supporting their potential for enhanced clinical applications. This study demonstrates that rational molecular design can improve both the optical properties and the drug-delivery proficiency of temoporfin, paving the way for more effective deep-tissue PDT treatments.

## Introduction

Although, it has been known since antiquity, the use of light in clinical applications has significantly evolved in modern practices. Phototherapy approaches correlate with the limitation of adverse side effects and, thus, globally improve the patients’ quality of life. Among the different flavors of phototherapy, photodynamic therapy (PDT) [1–4] deserves particular attention owing to its widespread use and efficiency.

PDT is a method of treatment taking advantage of the irradiation of a photosensitizer with a specific wavelength of light, which is then able to produce cytotoxic species, such as singlet oxygen, ^1^O_2_, which in turn induce cell death [2,5,6]. Light activated drugs offer a non-invasive and more localized treatment option compared to alternatives, such as chemotherapy. In PDT the PS is usually administered directly and systemically into the patient’s bloodstream [7]. Then the target tumorous area is irradiated with light of a specific wavelength enabling the excitation of the PS molecule to its singlet excited state manifold. The favorable route to generate singlet oxygen involves the long-lived population of the first triplet excited state (T_1_) via intersystem crossing (ISC) [8,9]. Persistent population of the triplet state allows the spin-allowed energy transfer with the ground state triplet oxygen, ^3^O_2_, resulting in the production of ^1^O_2_ and the relaxation of the PS to its singlet ground state. As the transition from T_1_ → S_0_ is a spin-forbidden process, the T_1_ states are usually longer-lived, and this time interval is enough for the photosensitizer to transfer its energy to the molecular oxygen (^3^O_2_) in its proximity. The general mechanism behind the PDT process is illustrated in Figure 1.

**Figure 1.**
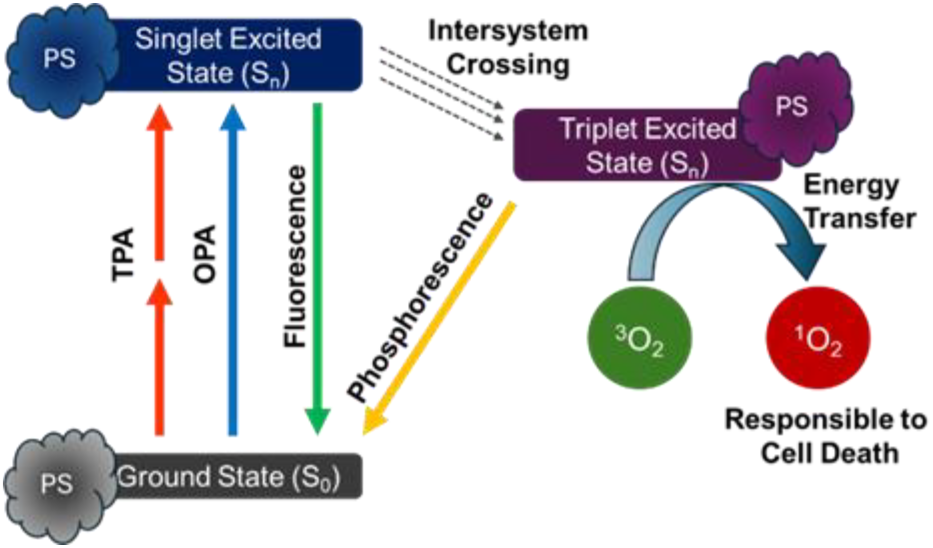
Mechanism of PDT action.

Even though PDT offers a non-invasive and sustainable process, its main challenges are linked to deep tissue penetration and selective targeting of cancer cells. Conceptually, an ideal PS should be delivered to the target tumor cells to increase treatment selectivity and also to reduce side-effects [10]. In fact, the coupling of PS with ligands targeting receptors overexpressed in cancer cells, such as folate acid, has been of particular interest recently [11–13]. While visible light can only be used on superficial or accessible lesions, due to its limited penetration, the development of PS absorbing in the near infared, and specifically in the therapeutic windows to which biological tissues are transparent, is highly suitable. Therefore, different aspects of PS should be considered concerning their optimization: their bioavailability, and ideally their ability to localize in tumor cells, their optical properties, and their photophysical evolution, which should allow efficient ISC and the subsequent generation of ROS. Presently, several PS have been clinically approved for PDT [14– 16]. However, the development of a PS presenting high tumor specificity, low toxicity in the dark, high cytotoxicity upon infrared light activation, sufficient lipophilicity to penetrate the cell membrane, and enhanced solubility in physiological environment remains challenging [17]. As a rule of thumb, a PS should present a good balance between hydrophilicity and lipophilicity to achieve both physiological solubility and membrane penetration. Temoporfin, also known as mTHPC (*meso*-Tetrahydroxyphenylchlorin), is a synthetic tetrapyrrole derivative that contains four phenol groups attached at the *meso* positions to form a chlorin cycle [18–20]. mTHPC has been used as a PDT agent under the commercial name Foscan^©^ since it was approved by the European Medicines Agency (EMA) in 2003. mTHPC is a second-generation photosensitizer, which presents a narrow and intense absorption band at 652 nm falling at the border of the clinical window, undergoes ISC with a high quantum yield, and is stable in biological media [18,19]. However, despite its promising features, the usage of mTHPC within the PDT framework is limited to topical applications or palliative treatments, especially for head and neck cancers, due to its tendency for aggregation, which limits its bioavailability and may alter its photophysical and optical properties. Even though mTHPC has already been approved and commercialized, several strategies are constantly being explored to overcome these shortcomings [18].

These include the encapsulation of mTHPC with a vectorization agent such as peptides [21], cyclodextrins (CDs) [22–25], and liposomes [26,27] to increase its solubility, targeting, and tissue penetration abilities. We have shown that the encapsulation of mTHPC can be readily achieved with β-cyclodextrin units in 2:1 ratio [28], without altering the ISC quantum yields of mTHPC [24], yet allowing the fast internalization of the drug-delivery complex and its possible spontaneous dissociation in the lipid bilayer core [23].

Other molecular design strategies are aimed at enhancing the infrared absorption properties of mTHPC to be used in deep tumor treatment. A possible strategy to tackle this issue is to rely on Two-Photon Absorbing (TPA) photosensitizers, which in this case may be introduced by suitable functionalization of the tetrapyrrole unit. TPA has been firstly theorized by Maria Göppert-Mayer in 1931 and later observed in the 1960s after the invention of lasers [29]. The mechanism involves the simultaneous absorption of two photons of equal or different energies, going through a virtual state to populate a suitable excited state. If we consider monochromatic light, the electronic transition can, thus, be achieved with photons of half the energy difference between the ground and the excited state, hence TPA takes places at wavelengths which are doubled compared to one photon absorption (OPA). However, since it is a formally forbidden process, TPA probability is usually smaller than single photon absorption. Furthermore, because of the need of simultaneous absorption of two photons, the absortion will depend on the square of the intensity of the light source. Yet, in addition to the displacement of the absorption towards longer wavelengths allowing deeper tissues penetration, TPA offers several advantages to improve PDT efficiency: the TPA cross-section quadratic dependence on the laser intensity increases spatial selectivity since the PDT activation will be restricted to the light-source focal point only minimizing lesions at the border, on the other hand, the low energy of the TPA wavelength minimizes adjacent tissue damage. Yet, such enhancements depend on the development of new photosensitizers characterized by high TPA cross sections (σ_TPA_) in the near infrared region, since those currently proposed still have low PDT efficiency.

In previous contributions, the σ_TPA_ for mTHPC has been estimated at 28 ± 8 GM at 775 nm in DMSO [30] and 18 GM at 800 nm in 20% ethanol, 30% polyethylene glycol and 50% distilled water [31]. While both experiments suggest the possibility of using mTHPC in TPA-based PDT, the cross section should be higher to allow for efficient light harvesting.

For this purpose, Senge *et al*. published a study in 2021 reporting the functionalization of mTHPC with different functional groups to systematically increase the σ_TPA_ while correlating the increase of the TPA cross-section with the PS structure [32].

From literature, it appears that the increase of σ_TPA_ in tetrapyrrolic systems can be achieved by different strategies. These include, the increasing of the conjugation length, the integration of ethynyl bridge at β [33] or *meso* position [34,35], and the functionalization with antennas that already possess a high σ_TPA_ [36]. Herein, we explore the modification of mTHPC with 3 different TPA-active dyes: DTP1, DTP2, and DPP (Figure 2) to increase TPA cross-sections. To the best of our knowledge, DTP derivatives have not been previously incorporated to improve the optical properties of tetrapyrrole-derived molecules. DTP are derived from the substitution of a π-chain and an acceptor group to different positions on the 2,5-dithienylpyrrole core [37–39], and have been extensively analyzed *via* experimental [37] and computational [38,39] methods. DTP1 showed two absorption peaks at 566 nm and 820 nm, whose TPA cross sections are found to be 422.0 and 1243.6 GM, respectively. DTP2, on the other hand, shows σ_TPA_ of 742 and 6977.7 GM for its peaks at 624 nm and 870 nm, respectively. Moreover, the carboxylic unit on the DTP moiety can be exploited to conjugate small molecules to target specific oncogenic receptors overexpressed in tumor cells [40,41]. Conversely, the TPA cross-section of polyethylene glycolated (PEG) diketopyrrollopyrrole (DPP) conjugated with a zinc-bound porphyrin (ZnP) reveals an elevation of the σ_TPA_ at λ_max_ from 20 GM to 1400 GM [35]. Therefore, the PEGylated DPP molecule has been included in this study since it provides a suitable comparison to unravel the properties of previously unstudied DTP antennas. However, to our knowledge, this specific temoporfin moiety has not been previously conjugated with PEGylated DPP. The main objective of this work is to improve the optical properties of mTHPC in order to be used as a PDT agent in the treatment of deep-tissue lesions. To this end, we have designed three novel molecules by combining three high σ_TPA_ compounds (DTP1, DTP2 and DPP) with the original mTHPC molecule (Figure 2).

**Figure 2.**
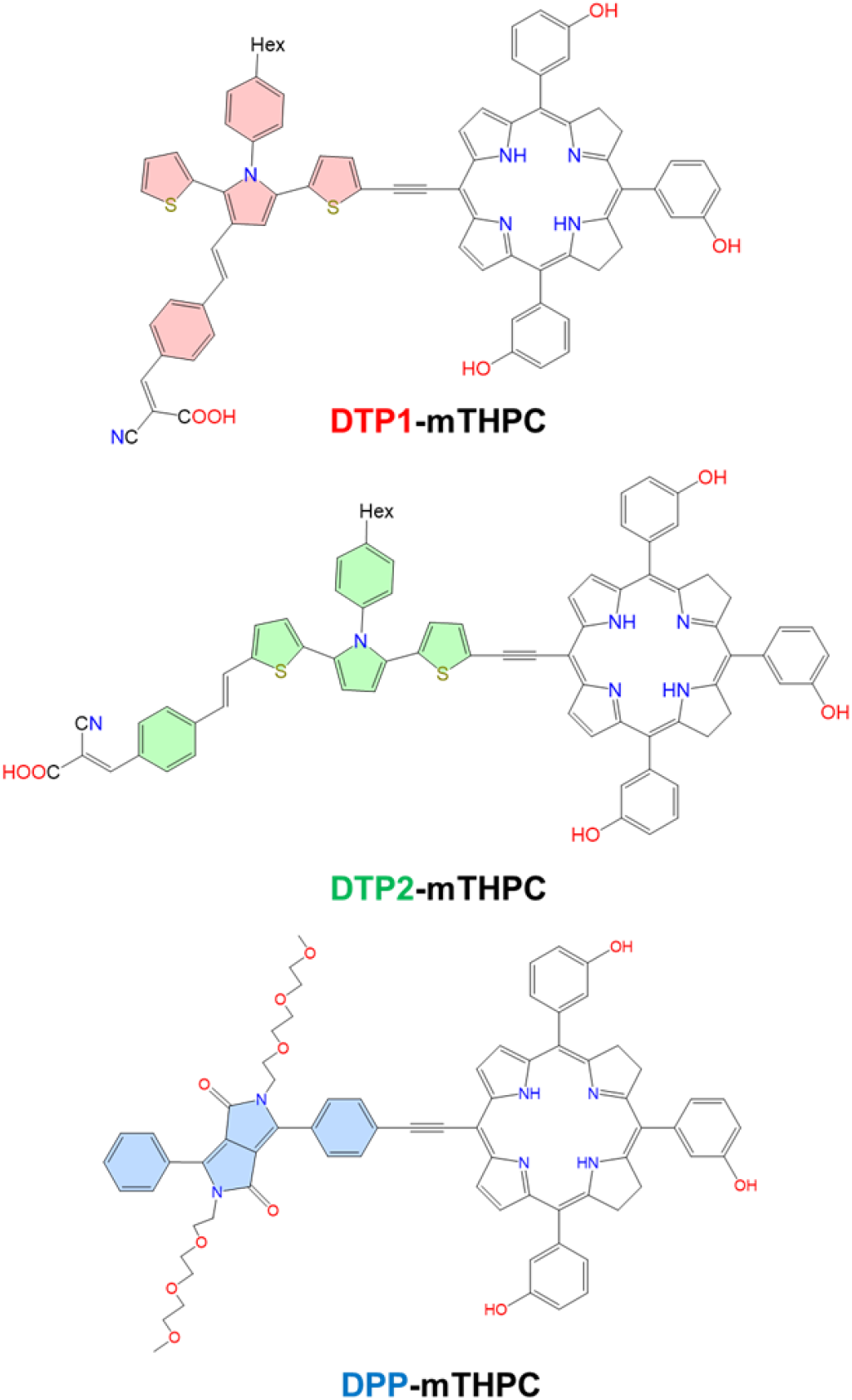
Structures of proposed TPA-sensitized photosensitizers, TPA antenna moieties are highlighted.

In line with this purpose, a variety of quantum mechanical (QM) analysis were used to predict the optical and photophysical properties of DTP- and DPP-conjugated mTHPC molecules, such as OPA and TPA cross sections, spin-orbit coupling (SOC), singlet and triplet energy gaps (Δ_ST_), and natural transition orbitals (NTO). Subsequently, benefiting from molecular dynamics (MD) simulations, PS were simulated in various environments that mimic biological conditions, such as bulk water and in interaction with a lipid bilayer representing a cellular membrane. Furthermore, to address the solubility issue, the candidate PS were encapsulated with β-CDs. Two CD units have been used to encapsulate DTP1-mTHPC and DTP2-mTHPC, while in the case of DPP-mTHPC only one CD was used since the presence of the two polyethylene glycol (PEG) groups induces potential steric clashes and is already improving the overall solubility. Randomly selected snapshots from classical MD simulations have also been extracted to calculate vertical absorption energies and cross-sections at the QM/MM level to observe if the absorption patterns were influenced by these environmental parameters.

## Computational Methodology

### Quantum Mechanical Calculations

All geometry optimization and single point excited states calculations were performed with the Gaussian 16 program package.[42] The ground and excited state geometry optimizations were done both *in vacuo* and in water solution modeled with Polarizable Continuum Method (PCM) [43,44]. Ground state geometry optimizations were performed at density functional theory (DFT) level using the meta-hybrid M06-2X [45] functional as its performance was shown to be superior in describing the ground state of chlorin type molecules compared to the B3LYP functional [46,47]. Though 6-31+G(d) basis set was shown to be sufficient to describe the electronic properties of mTHPC in a previous study [47], to better account for the changes rising from the inclusion of the TPA antenna the 6-31+G(d,p) basis set was employed for all QM calculations. On top of the optimized ground state geometry, the performance of different functionals in reproducing the vertical excitation energies using Time Dependent-DFT (TD-DFT) has been assessed. The selected exchange-correlation functionals included CAM-B3LYP [48], LC-BLYP[49] and ωB97XD [50] *in vacuo* and water and the results are reported in Supporting Information, Figure S1. To better compare with the experimental spectra all the vertical transitions have been convoluted using Gaussian functions of full-width at half length (FWHL) of 0.2 eV.

Furthermore, to obtain more realistic absorption properties, sampling of the Wigner distribution over the ground state geometry according to vibrational degrees of freedom and vibrational frequencies was performed via NewtonX program [51]. From 100 initial geometries, 40 structures were selected to perform TD-DFT calculations using the 3 functionals previously listed for the first 20 excited singlet states. Based on these results we selected ωB97XD functional with 6-31+G(d,p) basis set for further modeling and geometry optimization of the relevant excited states. This also aligns with the fact that ωB97XD functional has been shown to accurately describe the experimental vertical transitions of mTHPC and DTP derivatives [39,47]. To calculate σ_TPA_ of the antenna-mTHPC systems, the publicly available Dalton2020 program package [52] was used. The linear response methodology calculations were executed *in vacuo* at the CAM-B3LYP/6-31G(d) level of theory. In previous benchmarks from different studies, comparing different DFT functionals available in the Dalton package, CAM-B3LYP was found to outperform for TPA calculations [53,54]. Nancy_EX code [55–57] was used to obtain the natural transition orbitals (NTO’s) and the topological Φs indices, characterizing the electronic excited state’s nature in terms of electronic density rearrangement. NTOs are visualized with the Avogadro visualization and analysis program [58]. Additionally, Amsterdam Modeling Suite AMS/2022.103 package [59] is used for the calculation of the singlet-triplet gap and SOC at the DFT level with ωB97XD functional using 40 initial geometries sampled by Wigner distribution around the S_0_ and S_1_ minima, respectively. DZP basis set together with ZORA scalar and TD-DFT method was employed together within a water environment, which was modeled with the integral equation formalism polarizable continuum model (IEF-PCM). As ADF does not support Pople type basis sets, DZP basis has been chosen as a good compromise between accuracy and computational cost.

### Molecular Dynamics Simulations and QM/MM

The Amber22 program package [60,61] was used to perform classical MD calculations. Force field parameters for each candidate PS were optimized using the Antechamber package [62] in conjunction with Generalized Amber Force Field 2 (GAFF2) [63]. The structure of the CD units were obtained from the protein data bank (PDB ID:3CGT [64]) consistent with a previous study [65] and modeled using GLYCAM06_j [66] force field. The lipid bilayer used to model the biological membrane was built using the CHARMM-GUI[67] web-server and consists of 150 1-palmitoyl-2-oleoyl-sn-glycero-3-phosphocholine (POPC) units on each layer hydrated by 14808 TIP3P [68] water molecules, counterions (K^+^ and Cl^-^) are used to ensure a neutral simulation box and a physiological (0.15 M) salt concentration. POPC was chosen as the model lipid to allow a direct comparison with our previous work [23,47], and due to its ability to reproduce the behavior of eukaryotic cellular membranes, Amber Lipid17 Force Field [69] was used to represent the bilayer. It’s worth noting that in a previous study where we modeled mTHPC in similar environments, these computational conditions have proven capable of reproducing the experimental behavior [24].

Simulations were propagated in the constant pressure and temperature (NPT) ensemble at 1 atm and 300 K. Temperature control relied on the Langevin thermostat [70], while pressure regulation used the Berendsen barostat [71]. Long-range electrostatic interactions were calculated through the Particle-Mesh-Ewald method [72], employing an 8 Å cutoff distance, while periodic boundary condition (PBC) were consistently used. For each simulation, hydrogen mass repartition (HMR) [73] was used in combination with the Rattle and Shake algorithms [74] to slow the highest frequency vibrations involving hydrogens, thus allowing the use of a 4.0 fs time step to integrate the Newton’s equations of motion.

As aforementioned, three distinct environments were explored: aqueous phase, β-CD encapsulation, and interaction with a lipid bilayer. Equilibration involved gradual heating over 10 ps from 0 to 300 K. Membrane equilibration utilized a protocol for gradual restraint removal to ensure stability, therefore, the constraints on the heavy atoms of the POPC were progressively reduced in three consecutive steps of 36 ns each. The production simulations lasted 500 ns for each environment.

MD trajectories were analyzed with CPPTRAJ [75] of AmberTools23 [76] and VMD [77], the membrane thickness, area per lipid, and density profiles were obtained using the MEMBPLUGIN [78] extension and Density Profile Tool [79], respectively.

For the QM/MM Study of the absorption spectra, the Terachem/Amber [80] interface was used to obtain the absorption spectra of the PS in different simulated environments. Using the CPPTRAJ tool, 100 snapshots from each simulation except *in vacuo* have been extracted. On top of these snapshots vertical transitions have been calculated at the TD-DFT level with the ωB97XD/6-31+G(d,p) protocol. The whole TPA-mTHPC was included in the quantum partition, whereas water, β-CD and POPC lipids are treated at MM level.

## Results and Discussion

### Optical Properties

The simulated OPA and TPA absorption spectrum for the three antenna-sensitized chromophores are reported in Figure 3, together with the most relevant NTOs describing the lowest state with highest cross-section for OPA and TPA, respectively. As expected, the shape of the OPA spectrum resembles one of a porphyrin-like systems, notably characterized by the presence of the Q- and Soret-bands in the visible and infrared region, respectively. Notably, the position of the Q-band is only slightly altered by the presence of the antenna. In addition, and mostly in the visible range, additional bands which may be assigned to absorption centered on the antennas are also present. Intermolecular charge-transfer states featuring transitions between the two moieties are safely excluded, at least in the low energy portion of the spectrum. However, especially in the case of the DTP2-sensitized compound delocalized excitations spanning the whole molecular systems can be observed, notably for S_4_. The fluorescence absorption (Figure S6) has also been modeled showing that for each molecule the emission takes place from a porphyrin-centered state, and thus is only marginally sensitive to the presence of the antenna.

**Figure 3.**
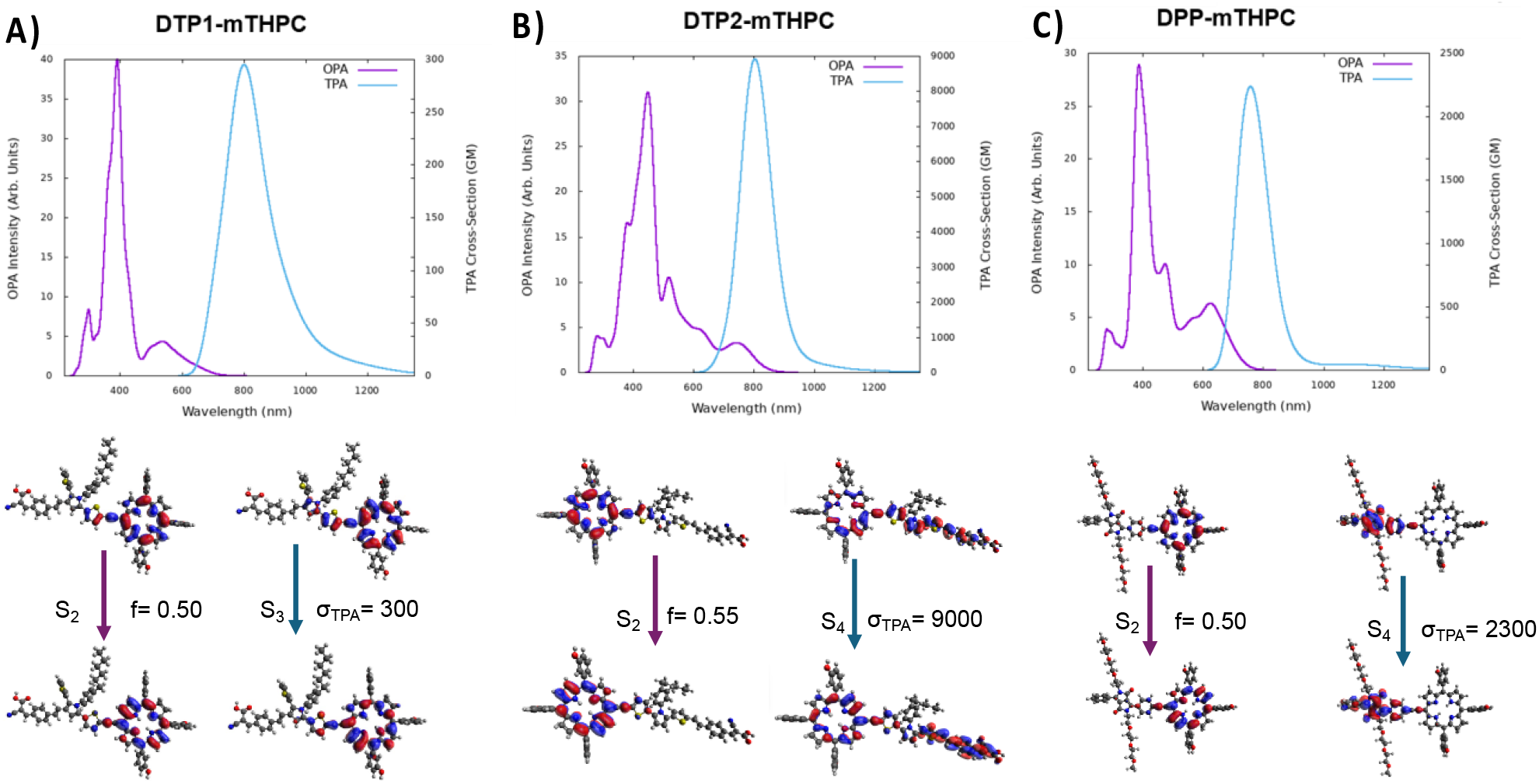
Normalized One-photon (ωB97XD/6-31+G(d,p)) and Two-photon (CAM-B3LYP/6-31G(d)) absorption spectra of a) DTP1-mTHPC b) DTP2-mTHPC and c) DPP-mTHPC molecules, together with their corresponding λ_max_ (eV) and σ (GM) values.

However, if the modification of the one-photon optical properties brought by the antenna are only marginal, more subtle and significant changes can be accessed in the TPA spectra. Indeed, it was reported that the functionalization of the parent chromophore such as through the addition of carbonyl groups, alkyne linkers and high two-photon absorbing molecules have a significant effect in increasing the TPA cross-section [81–86].

All three antenna-sensitized molecules show a relevant and rather intense TPA signal in the infared region (Figure 3), however significant differences should be pointed out. Indeed, mTHPC-DTP1 presents a large and rather asymmetric band. Interestingly, the most intense TPA excitation involves the porphyrin Q-band S_3_ state peaking at 300 GM. The asymmetry of the band is instead due to the TPA excitation of the DTP1 moiety, which has, however a lower inherent cross-section as already shown in our previous work. Therefore, the functionalization with DTP1 appears suboptimal for the purpose of increasing the TPA capacity of the PS. This situation is instead reverted in the case of mTHPC-DPP, which presents a high cross-section at around 800 nm exceeding 2250 GM.

Interestingly, as shown by the corresponding NTOs, this transition is due to a DPP-based excited state, which is inherently TPA-active. However, the most striking TPA potential is observed for mTHPC-DTP2, which presents an impressive infrared cross-section achieving 9000 GM. This value largely outperforms the performance observed for the parent DTP2, which already presented the best TPA cross-section among its series. The enhancement of TPA activity upon conjugation with the porphyrin core may be rationalized by the presence of a strongly delocalized excitation (S4), which extends over the whole molecular system. This result is also extremely promising for potential applications, since mTHP-DTP2 presents a strong TPA in a spectral region compatible with the therapeutic window, thus allowing the treatment of deeper lesions.

### Photophysics and Intersystem Crossing Efficiency

The presence of a facile intersystem crossing (ISC) presenting a high quantum yield is fundamental to allow the population of the triplet state and, consequently, the energy transfer to the ground state ^3^O_2_ leading to the activation of the cytotoxic ^1^O_2_. To analyze the suitability of the ISC relaxation channel we have obtained the energy level diagram of the singlet and triplet states at Franck-Condon (SI) and at the S_1_ optimized geometry (Figure 4). Indeed, a region of singlet-triplet quasi-degeneration is needed to allow the non-adiabatic ISC process. Furthermore, since an ultrafast relaxation via the fully spin-allowed internal conversion (IC) is expected the topology of the potential energy surfaces at the S_1_ equilibrium geometry is the most crucial one.

**Figure 4.**
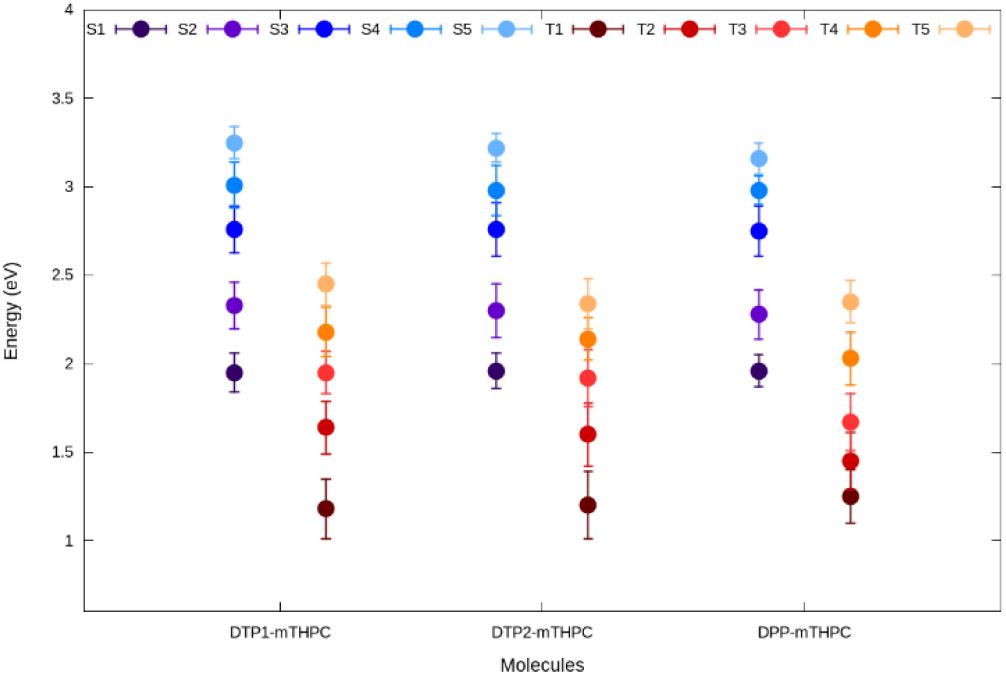
Average value of the singlet and triplet energy obtained by sampling the S_1_ minimum via a Wigner distribution. The standard deviation is also given

In Figure 4 we report the average energy value of the singlet and triplet states obtained from a Wigner sampling of the S_1_ minimum. The standard deviation is also provided to highlight the spread of the energy distribution upon vibrational movement.

We may immediately observe that mTHPC-DTP1 and mTHPC-DTP2 present nearly ideal conditions to assure ISC. Indeed a (quasi)-degeneracy between the energy of the S1 and the T3 state exist for both systems, the maximum gap observed for mTHPC-DTP2 being of only 0.038 eV. Therefore, we may hypothesize that both systems will follow a photophysical pathway involving the S_1_→T_3_ ISC followed by fast IC in the triplet manifold to reach T_1_. While this pathway is probable, given the energy degeneracy, it should be pointed out that SOC elements between S_1_ and T_3_ are weak amounting at 0.136 and 0.142 cm^-1^ for mTHPC-DTP1 and mTHPC-DTP2, respectively. Even if the low value of the SOC can significantly slow ISC, similar coupling elements are observed for the parent mTHPC, which is known to activate singlet oxygen, confirming the suitability of the proposed approach.

Conversely, mTHPC-DPP stands apart showing larger differences in the distribution of the excited states. Indeed, for this molecule the T4 average energy level lies 0.068 eV higher than S1, while T3 is at a distance of about 0.25 eV from the first singlet excited state. However, since the distribution of the energy levels of T4 and S1 sampled by a Wigner distribution, are still largely overlapping (Figure 4), we may thus hypothesize that the ISC will proceed through the non-adiabatic transition to T4, probably characterized by a lower quantum yield compared to the other chromophores, followed once again by the fast IC to reach the population of T_1_. As it was the case for the other compounds the SOC elements between the two most plausible states are weak (0.267 cm^-1^) further pointing to the non-ultrafast and non-unitary quantum yield of the process.

### Interaction of PS with Lipid Bilayer and Passive Drug Delivery

Having confirmed that the proposed PS candidates may be regarded as efficient TPA sensitizers, which can potentially undergo ISC and, hence, singlet oxygen activation, we decided to study their interaction with a model lipid bilayer which represents an ideal target for the PDT approach. In a previous study [24], we have already shown that the parent mTHPC is internalized in the lipid bilayer residing in proximity to the polar heads, its density distribution partially overlapping with the position of the lipid tail double-bond, which is a target of oxidation by ^1^O_2_. By using classical MD simulations, we have shown (Figure 5) that all candidate PS molecules interact persistently with the lipid bilayer and in particular position themselves in the polar head-region. The formation of the membrane bound molecule takes place spontaneously and is persistent along the time-scale of the MD simulation suggesting the formation of a stable complex (Figure S9 and S10). Consistent with the behavior of the parent porphyrin moiety we observe a partial overlapping of the density distribution of the PS with that of the lipid tail, indicating a favorable positioning to allow ^1^O_2_ activation in proximity to its biological target. Interestingly, mTHPC-DPP seems to penetrate deeper into the membrane hydrophobic core as shown by its density profile. However, this fact is mostly due to the PEG tails rather than the porphyrin core which remains strongly anchored to the polar head region.

**Figure 5.**
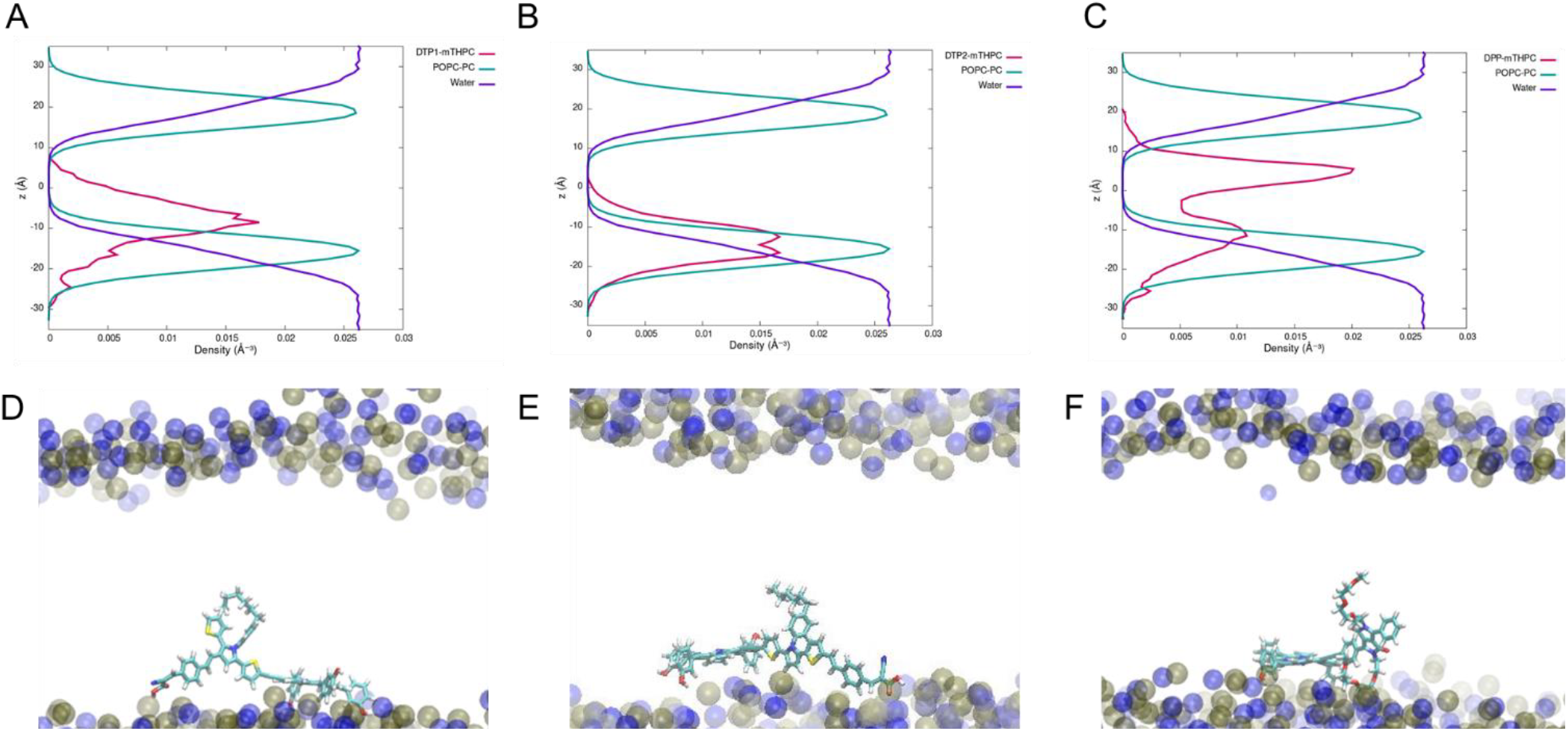
Density distribution profile along the membrane axis for the chromophore, the lipid tails, and water for mTHPC-DTP1 (A), mTHPC-DTP2 (B), and mTHPC-DPP (C). Representative snapshots at the equilibrium position are shown for mTHPC-DTP1 (D), mTHPC-DTP2 (E), and mTHPC-DPP (F).

As shown in the SI (Figures S11-S13) the chromophore is maintained at the polar head interface by a subtle combination and balance of different interactions involving notably dispersion interactions between the lipid tails and the hydrophobic core of the chromophore. In addition, cation-π and alkyl-π interactions are formed with the phosphatidyl choline, palmitoyl acyl and monounsaturated oleoyl acyl moieties (Tables S15-S17). Interestingly, in the case of mTHPC-DPP only the porphyrin core is interacting with the polar head, leaving the antenna free to bury into the lipid bilayer. Interestingly, mTHPC is also the only compound showing the emergence of hydrogen bonds albeit quite labile.

One of the main drawbacks suffered by PDT drugs is their limited bioavailability, due to the presence of extended conjugated moieties leading to poor water solubility. To overcome this issue passive drug delivery strategies can be envisaged consisting of encapsulating the drug with a water-soluble hollow carrier, such as cyclodextrin (CD) [28] or calixarenes[87]. In previous studies [23,24], we have shown that mTPHC encapsulation by β-CD is an efficient strategy increasing water solubility without preventing the internalization in the lipid membrane. Thus, we explore this possibility with our proposed PS anchored with TPA antenna. As seen in Figure 6 and Figure S14, the DTP1- and DTP2-functionalized chromophores form a 1:2 complex with β-CD, similar to the behavior of the parent mTHPC. Notably while the β-CD is encapsulating the porphyrin core, the TPA antenna remains largely exposed to the solvent. This fact can, however, facilitate its inclusion in the lipid bilayer enhancing the amphiphilicity of the drug. Both DTP1 and DTP2 give rise to stable and persistent complexes and the distance between the center of mass of mTHPC and the β-CD remains stable with a distribution peaked at around 8 Å. Interestingly, DTP2 seems to form a slightly more loosely bound complex presenting a larger distance between the center of mass at about 9 Å. In the case of mTHPC-DPP we only attempted to build a 1:1 complex with CD, due to the larger steric hindrance of the PEG moieties, and their increased hydrophilic character. However, the complex with the CD is highly persistent even if the maximum value of the distance between the center of mass increases up to 11 Å. Similar to what was observed for the parent mTHPC, the encapsulation reduces the rotational movements of the phenyl substituents, while the flexibility of the solvent exposed antenna is not altered (Figure S15).

**Figure 6.**
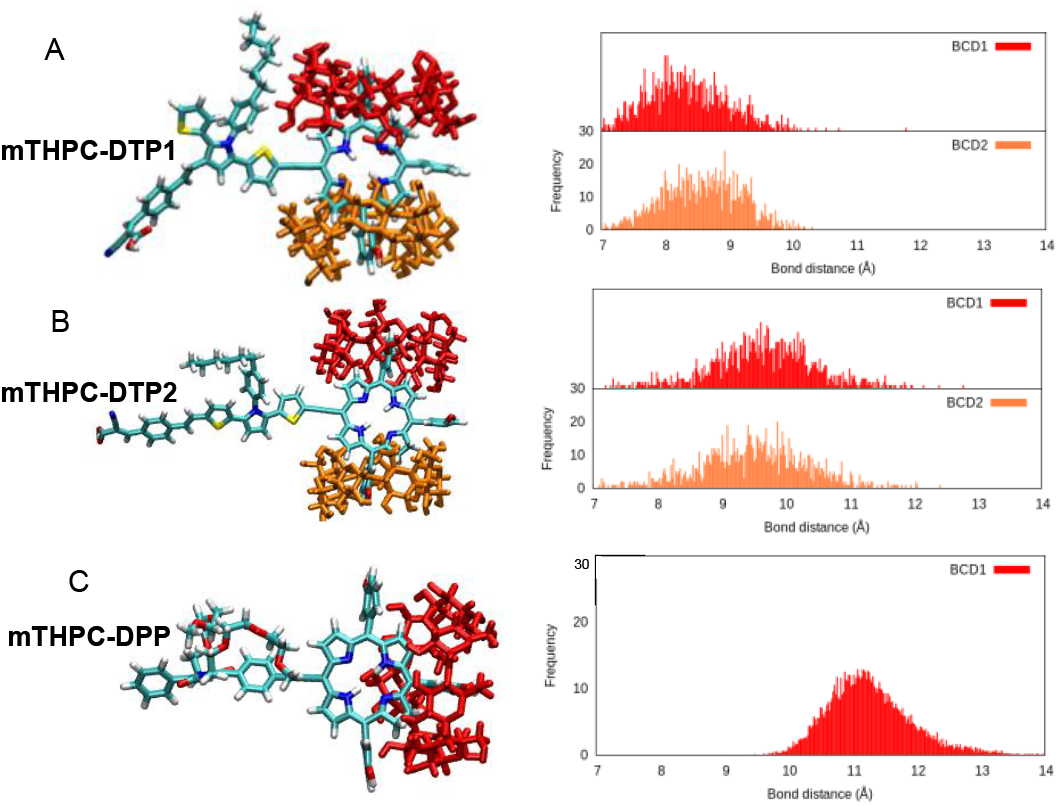
Representative snapshots of the complex between the PS and CD, the distribution of the distance between the center of mass of mTHPC and the complexing CD is also reported. A) mTHPC-DTP1, B) mTHPC-DTP2, C) mTHPC-DPP.

Finally, to assess the influence of the encapsulation and membrane interaction on the optical properties of our chromophores we calculated the OPA spectrum on top of snapshots extracted from MD with the chromophore placed either in water, inside a lipid bilayer, or encapsulated with CD (Figure 7). Besides some minor differences our results confirm that the main features of the spectra as described in the previous section are largely unchanged. More importantly, no significant modification in the position or the shape of the main absorption bands is observed between the different environments. This result is highly encouraging since it suggests that the photophysical pathway leading favorably to ^1^O_2_ activation should hold under different conditions potentially experienced by the PDT agent. However, a further study involving the explicit determination of the photophysical pathways of TPA-decorated mTHPC in different explicit environments should be performed, ideally with non-adiabatic QM/MM dynamics to confirm this assumption.

**Figure 7.**
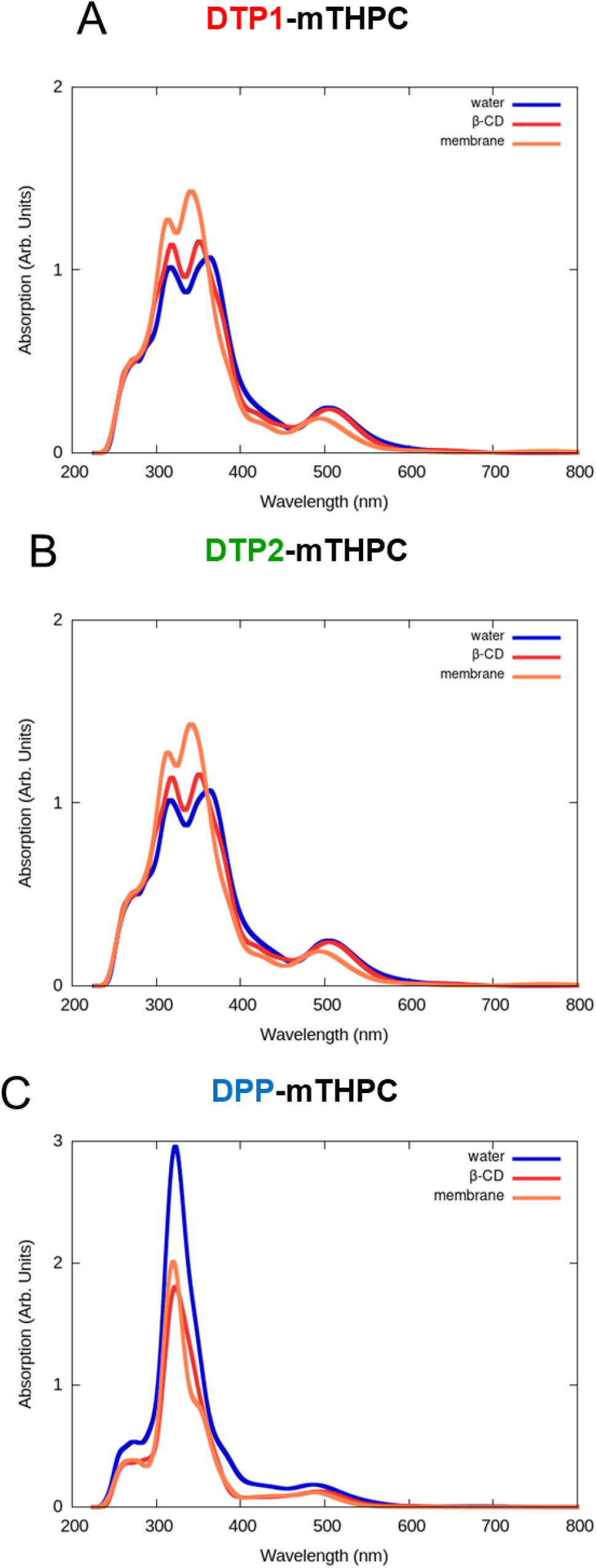
Calculated (at ωB97XD/6-31+G(d,p)) normalized absorption spectrum of TPA-anchored mTHPC molecules in different environments: in water (blue), 2PA-mTHPC:β-CD in water (red) and in proximity of the lipid membrane (orange). The ground state conformational space has been sampled with 500 ns equilibrium MD.

## Conclusion

By resorting to a multiscale approach involving ground and excited state quantum chemistry simulations as well as classical MD and QM/MM simulations, we proposed an original strategy to enhance the PDT efficiency of mTHPC drug. In particular, we show that the functionalization of mTHPC with different TPA antenna is suitable to significantly increase the TPA cross-section leading to an intense absorption band in the infrared region. In particular the functionalization with DTP2 leads to an impressive TPA cross-section σ_TPA_ of about 9000 GM, rarely observed for organic chromophores. Furthermore, we have also shown that the TPA functionalization preserve the favorable photophysical routes leading to ISC and, thus ^1^O_2_ activation. Indeed, especially in the case of DTP1- and DTP2-functionalized PS an almost perfect degeneracy between T_3_ and S1 holds at the S_1_ equilibrium geometry. Such alignment of states should provide efficient ISC, despite the small values of the SOC. On the contrary ISC for mTHPC-DPP should be slightly less efficient and be allowed by vibrational activation.

Finally, we have confirmed that all the TPA-functionalized compounds may be readily internalized in lipid bilayers, persistently residing at the polar head and presenting some overlap with the position of the lipid tail double bonds, which are the primary target of the activated singlet oxygen. In addition, we have also confirmed that the TPA-sensitized PS could potentially be encapsulated with CD moieties for passive delivery, consistent with the strategy used for the parent mTHPC drug. Importantly, the different molecular environments do not alter the optical properties of the drug candidates, suggesting the robustness of the ISC pathways.

Our computational approach has allowed the proposition of TPA-sensitized PS, which present extremely high TPA cross-sections combined with a facile ISC and membrane internalization capabilities. As such they may represent a most promising solution for the PDT treatment of deeper lesions exploiting IR excitation. In the future, we plan to study the interaction of the encapsulated PS with the model lipid bilayer to confirm the efficient delivery of the drug. Furthermore, the photophysical pathway leading to ISC upon excitation of the TPA-active singlet state will be explored by surface-hopping non-adiabatic dynamics simulations, eventually at the QM/MM level. Yet, we believe that our study already highlights the power of computational chemistry and rational molecular design in proposing novel PDT agents, paving the way for future experimental validation and clinical translation.

## Supporting Information

Ground Singlet State (S_0_) and first excited singlet state (S_1_) Geometries, QM functional performance, electronic structure calculations at QM level, PS Molecules within the biological environment including RMSD graphs, PS Molecules interacting with membrane components, and encapsulated PS Molecules with β-Cyclodextrin Units (file type, PDF). The authors have cited additional references within the Supporting Information [37– 39,47].

## Acknowledgements

The authors thank GENCI, Explor and National Center for High Performance Computing of Turkey (UHeM) under grant number 1011062021, computing centers and the Platform P3MB for computational resources. The authors thanks ANR and CGI for their financial support of this work through Labex SEAM ANR 11 LABEX 086, ANR 11 IDEX 05 02 and PIRATE. The support of the IdEx “Université Paris 2019” ANR-18-IDEX-0001. Support from the PEPR LUMA is also gratefully acknowledged. BKF and SC thank TUBITAK (Project Number: 120Z659) for financial support. BKF would also like to thank the French Embassy in Turkey for a joint PhD grant.

---

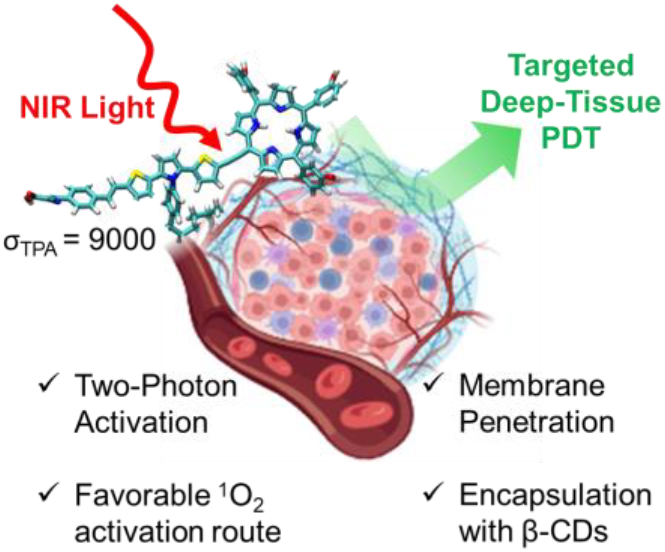
Entry for the Table of Contents Institute and/or researcher Twitter usernames: @AntonioMonari

